# A Quantitative Polymerase Chain Reaction Protocol for Sex Identification of Zebra Finch and Chicken Using Blood Samples

**DOI:** 10.1101/2025.08.27.671803

**Authors:** Prakrit Subba, Saidat O. Adeniran-Obey, Fanny-Linn Kraft, Susan C. Chapman, Natalie A. Shay, Shannon R. Liedl, Mark D. Wild, Scarlett A. Wolcott, Julia M. George

**Author notes:** Correspondence should be addressed to Julia M. George.

## Abstract

Studies of early development in birds typically rely on PCR analysis of genomic DNA to identify embryonic or neonatal sex. In zebra finches and other birds, males are the homogametic sex (ZZ) while females are heterogametic (ZW), and females are distinguished by the presence of specific sequences on the female-specific W chromosome. However, when only a single W locus is analyzed, lack of a PCR product in a sample could potentially arise from genetic variation or technical failure of the amplification, leading to false identification of female samples as males. To mitigate this concern, we developed an approach that targets two different W loci, using SYBR-based quantitative PCR to analyze amplification curves. We applied this method to determine sex of 30 zebra finch embryos (embryonic day 13) and subsequently confirmed genetic sex by brain transcriptome sequencing. We also identified and tested primer sets that are effective for sex determination in chickens.

## Introduction

The zebra finch (*Taeniopygia guttata*) is a well-established model for studying vocal learning, neuroplasticity, and neuro-endocrine regulation [1–3]. In this species, obvious differences in morphology and behavior do not begin to emerge until the juvenile period, yet differences in gene expression and growth rate can be detected as early as 36h of embryonic incubation [4]. Many other bird species (e.g. owls, crows, and doves) are monomorphic, meaning that the sexes are not visually distinguishable even as adults. For this reason, reliable methods for determining genetic sex are critical for studying and managing captive and wild bird populations [5–7] and ensuring sex balance in experimental design for developmental and genomic research [8–12].

The avian sex chromosomes Z and W determine sex in birds, wherein males are homogametic (ZZ), while females are heterogametic (ZW). Existing methods for molecular sexing employ genomic fragment amplification using polymerase chain reaction (PCR) or real-time PCR methods to amplify sex-linked homologs of the chromodomain helicase DNA binding protein 1 gene (CHD1W and CHD1Z) [5,13–18]. Amplification of these two genes with conserved primers gives rise to products of different lengths, due to differences in the length of an intron, so that males (ZZ) yield a single PCR product while females (ZW) yield two. However, this approach is vulnerable to false results due to underlying genetic polymorphisms in the CHD1 sequences [19,20]. To address this concern, we optimized a qPCR protocol for the molecular sexing of zebra finches with genomic DNA extracted from small volumes of blood, using one primer set targeting sequences on the Z chromosome, and two primer sets targeting sequences on the W chromosome. We further validated this approach with novel primers designed for chicken and applied to blood sampled from adults of known sex.

## Materials and Methods

### Blood Sample Collection from Zebra Finch Embryos

Zebra finch eggs were laid by parents in an aviary at Clemson University. Birds were housed in a free-flight aviary (147 sq ft) on a 12:12 light/dark cycle at 22 °C (±1°C) with ∼50% humidity and were provided with *ad libitum* commercial seed diet, water, cuttlefish, grit, and supplemented twice **we**ekly with fresh spinach and boiled eggs. Nests were checked regularly for eggs and the embryos were staged by reference to a zebra finch-specific staging guide [21]. On embryonic day 13 (E13), the eggshells were carefully opened, and embryos (n = 30) were rapidly killed by decapitation with a clean razor blade. 2-5 μL of blood found inside the eggshell or trunk blood (i.e., when eggshell had lower volumes of blood) was collected in 1.5 mL microcentrifuge tubes, flash-frozen and stored in a -80°C freezer until use. All procedures wer**e** approved by the Clemson University Institutional Animal Care and Use Committee (IACUC), under protocols AUP2022-0486 and AUP2023-0106. Genomic DNA was extracte**d** from blood samples using either Chelex-100 resin (Sigma-Aldrich, Product No. C7901-25G) [22] or a commercial gDNA isolation kit (Zymo Quick-DNA Microprep Kit; Cat #D3021). DNA quality and quantity were assessed using a Nanodrop spectrophotometer, and the samples were stored at -20°C until further use.

### Blood Sample Collection from Adult Zebra Finches of Known Sex

The adult zebra finches used in this study were originally part of separate studies and breeding experiments at Deakin University, Australia [23,24]. The blood samples were sourced opportunistically between breeding attempts. The adults were housed with a colony of other zebra finches in a 3×2×2 m room. They were housed at a 14:10 light/dark cycle at 20°C (±1°C) with 50% humidity and were provided with *ad libitum* commercial seed diet, water, cuttlefish, grit, and cucumber. When the birds were breeding, they also received daily supplemental food consisting of hardboiled eggs and spinach.

Blood samples were collected when the birds were not breeding (i.e., when moved from single-sex housing or after the breeding experiment was completed). The birds were captured and held in opaque cloth bags until the blood samples were taken. We collected each blood sample by puncturing the alar vein with a 26.5 g needle and collecting 50–75 μL of blood with heparinized micro-hematocrit tubes. The blood samples were collected under the ethics permit G15-2015, approved by the Animal Ethics Committee Laboratory-Geelong (AECL-G). DNA was isolated from clotted whole blood samples stored in 70% ethanol [25,26].

### Blood Sample Collection from Adult Chicken

Blood was collected from adult birds at the Clemson University Poultry Farm (IACUC protocol AUP2024-0092). A 21 g needle was used to pierce the brachial wing vein, and 50 μL collected using a syringe. Blood was stored in a sterile microcentrifuge tube at -80°C until further use. Genomic DNA was extracted from blood samples using a commercial gDNA isolation kit (Zymo Quick-DNA Microprep Kit; Cat #D3021).

### Zebra Finch qPCR Primer Design and Annealing Temperature Optimization

We designed qPCR primers targeting genes on the Z and W chromosomes from the zebra finch genome assembly bTaeGut1.4.pri, two genes on the W chromosome LOC116806872 (nipped-B like protein) and LOC116806878 (mothers against decapentaplegic homolog 4) and one Z chromosome gene, PIK3C3 (phosphatidylinositol 3-kinase catalytic subunit type 3) (Table 1). Primers were designed using NCBI Primer Blast, specifying input gene coordinates, amplicon size (100-250bp), primer melting temperature (minimum = 57.0°C, optimum = 60.0°C, maximum = 63.0°C), max T_m_ difference = 3, database “nr”, and taxid:59729 (zebra finch). Primers were ordered “LabReady” from IDT (100um in 10 mM Tris, 0.1 mM EDTA, pH 8.0). To find the optimal annealing temperature for all zebra finch primer pairs, we used the temperature gradient feature the Bio-Rad CFX96 Real-Time System. We found that an annealing temperature of 58.0°C gave a single peak by melt curve analysis, confirming target specificity.

**Table 1:**
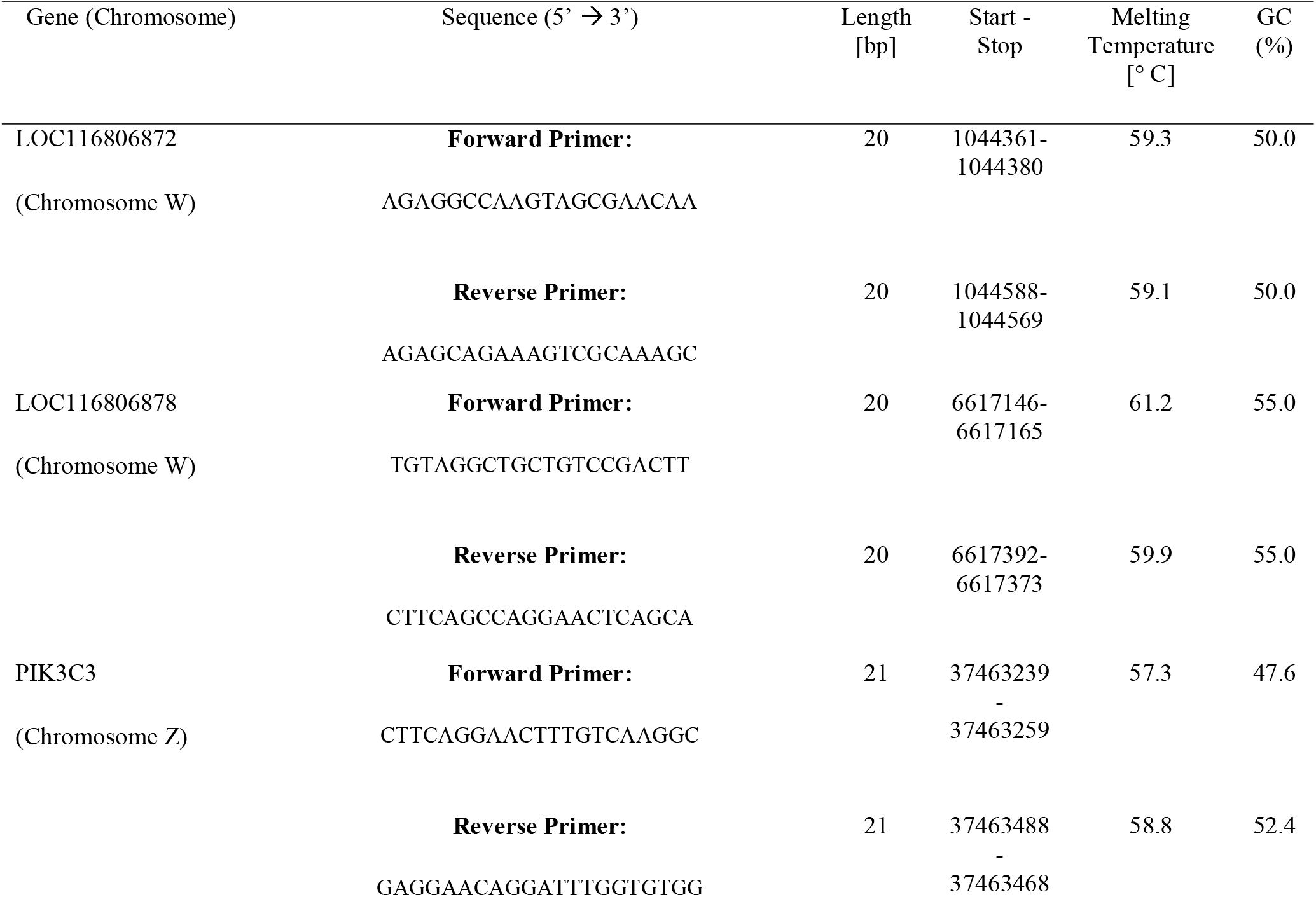
Primer Design for Zebra Finch Sex Chromosome Genes

### Chicken qPCR Primer Design and Annealing Temperature Optimization

To adapt our protocol in chicken, we selected the chicken orthologs of the Nipped-B homolog-like gene (LOC427025) on the W chromosome, the SPIN1L (spindlin 1 like) on the W chromosome, and PIK3C3 on the Z chromosome (Table 2), using the latest genome assembly (bGalGal1.mat.broiler.GRCg7b). Primers were designed using NCBI Primer Blast as above, and “*Gallus gallus* (taxid:9031)” was selected as the organism. We found that an annealing temperature of 61.0°C gave a single peak for each primer pair by melt curve analysis, confirming target specificity.

**Table 2:**
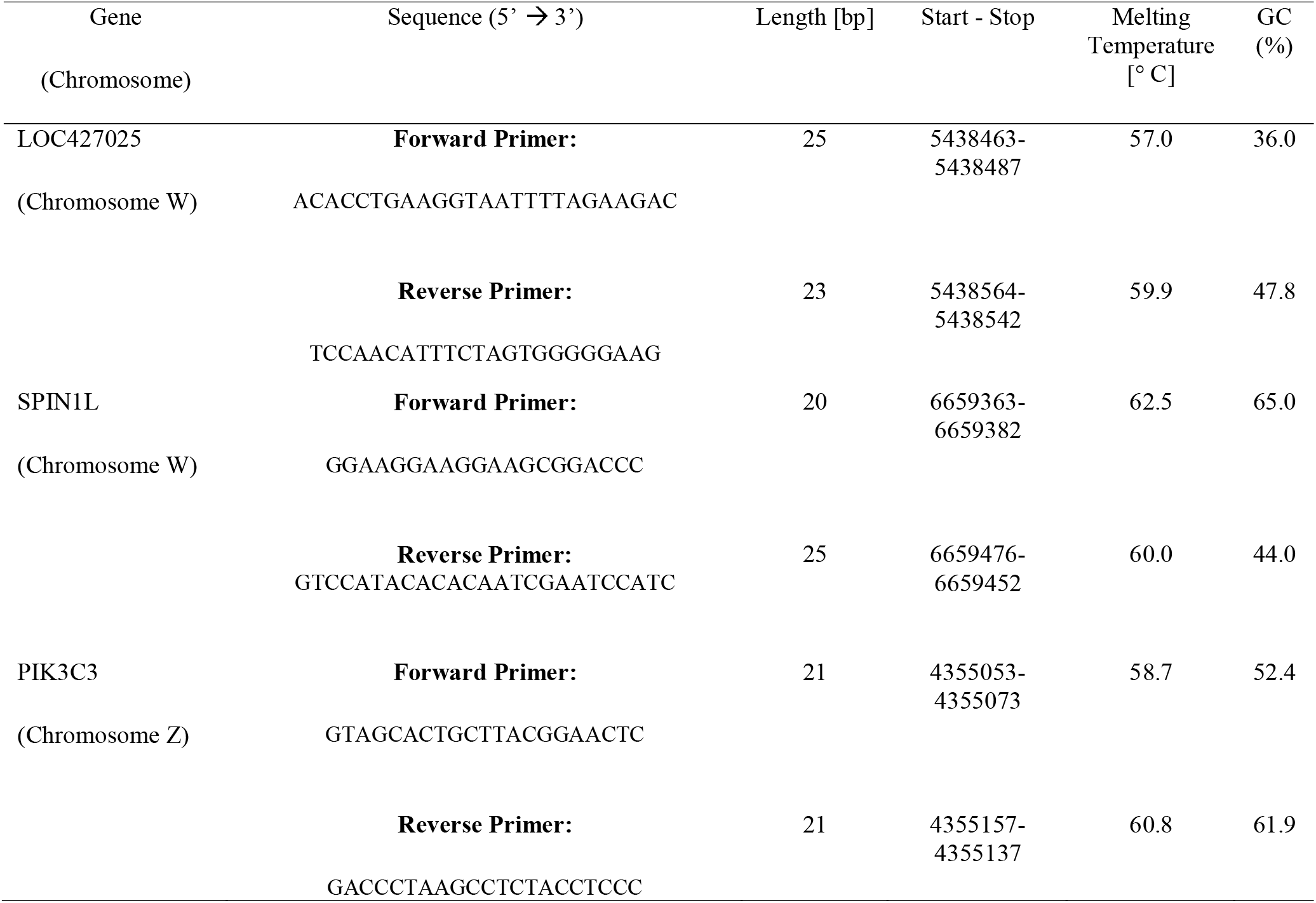
Primer Design for Chicken Sex Chromosome Genes.

### qPCR Reaction

Each 20 μL qPCR reaction contained 10 μL of 2x iTaq Universal SYBR Green Supermix (Bio-Rad Catalog #1725121), 2 μL each of forward and reverse primers (300–500□M final concentration per primer) and 6 μL of the gDNA product (1 ng gDNA) or non-template negative control. The total reaction volume was adjusted to 20 μL with nuclease-free water. All gDNA samples were run in triplicate for each primer pair in separate wells of a 96-well plate. Every plate included a negative control (deionized water instead of gDNA) and a positive control of a known male and female. The amplification and melt peak curves were then visualized using the Bio-Rad CFX Maestro software (version 2.2). The data files generated were downloaded in the CSV format and visualized using R (v 4.4.1).

### RNA Sequencing Validation of Zebra Finch Embryo Sex Identification

To validate the results of the qPCR-based sex identification, we performed RNA-sequencing on whole brains of E13 embryos (n = 30) that had been sexed by qPCR at collection. We extracted brain RNA using RNeasy Mini Kit (Cat. No. 74104) and sequenced stranded mRNA libraries on the NextSeq 2000 platform in the paired-end mode. Quality control was performed by trimming low quality sequences using Trimmomatic (v 0.39) [27]. The RNA-seq reads were mapped to the zebra finch reference transcriptome (bTaeGut1.4.pri) and the transcripts were quantified using Salmon (v 1.5.2) [28]. Counts were imported into DESeq2 (v 1.46.0) [29] to produce plots of normalized counts and conduct principal components analysis to highlight differences between identified males and females.

## Results

We compared two methods of blood gDNA extractions to find the most efficient protocol that provided the highest quality of gDNA. We extracted gDNA using either Chelex-100 resin or a gDNA preparation kit (Zymo Quick-DNA Micro Kit). The Zymo kit produced significantly higher A260/280 ratios (median = 1.885) compared to the Chelex gDNA method (median = 1.54; Mann-Whitney U = 15.5, p < 0.001). The rank-biserial correlation (r = -0.862) indicated a large effect size, with Zymo samples ranking consistently higher in DNA purity (Supplemental File 1). The gDNA purification kit was faster (< 1.5 hours) than the Chelex-100 protocol (9 hours).

To validate the protocol for the sex identification of zebra finches, SYBR-based qPCR was performed on DNA extracted from adult zebra finches of known sex. In males (ZZ), PIK3C3 primers produced an amplification curve while the W-specific targets were not detected (Figure 1a). The PIK3C3 primers amplified a single and specific genomic region, as evidenced by a single sharp peak upon melting curve analysis (Figure 1b). In females (ZW), the results show clear amplification and single melt curve peaks for PIK3C3 (Z chromosome), LOC116806872 (W chromosome) and LOC116806878 (W chromosome) (Figure 2a-b).

**Figure 1:**
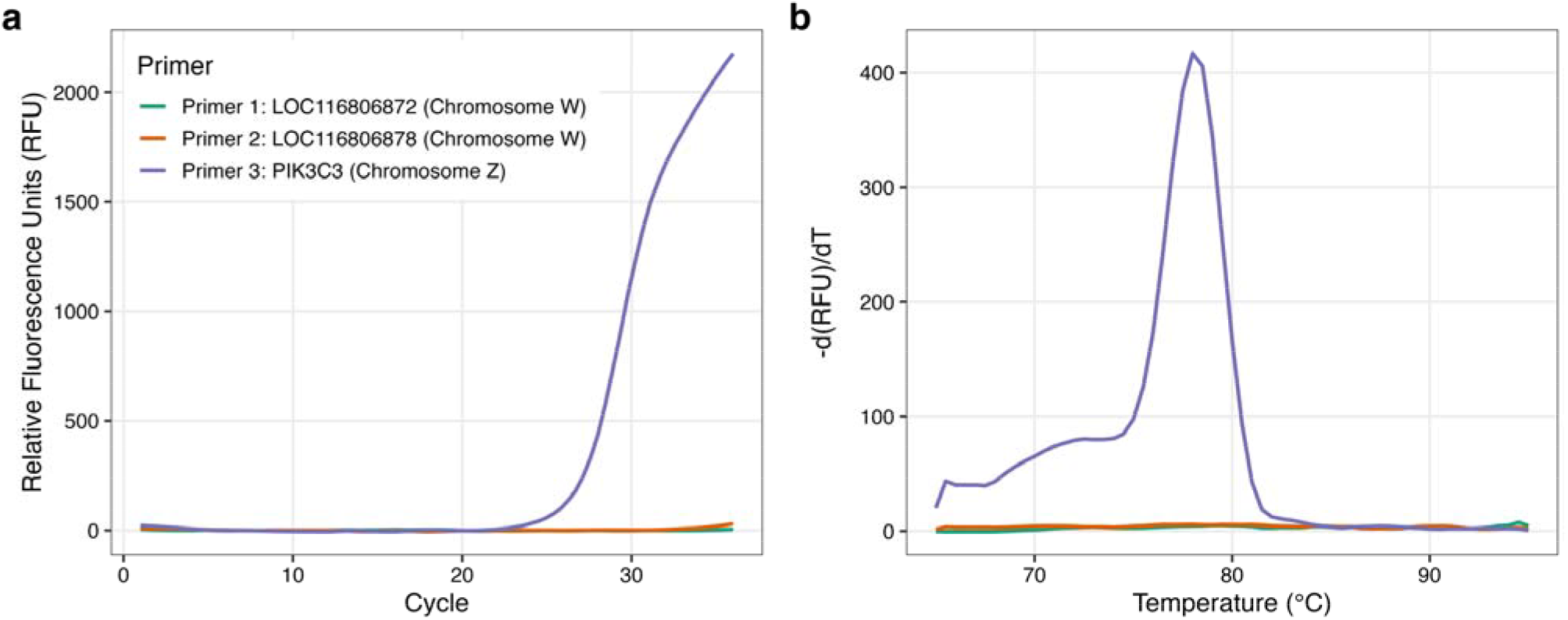
(a) DNA amplification (left) and (b) melting curve peak (right) from a known ZZ zebra finch adult male produces a single amplification curve for the Z chromosome-specific PIK3C3 gene.

**Figure 2:**
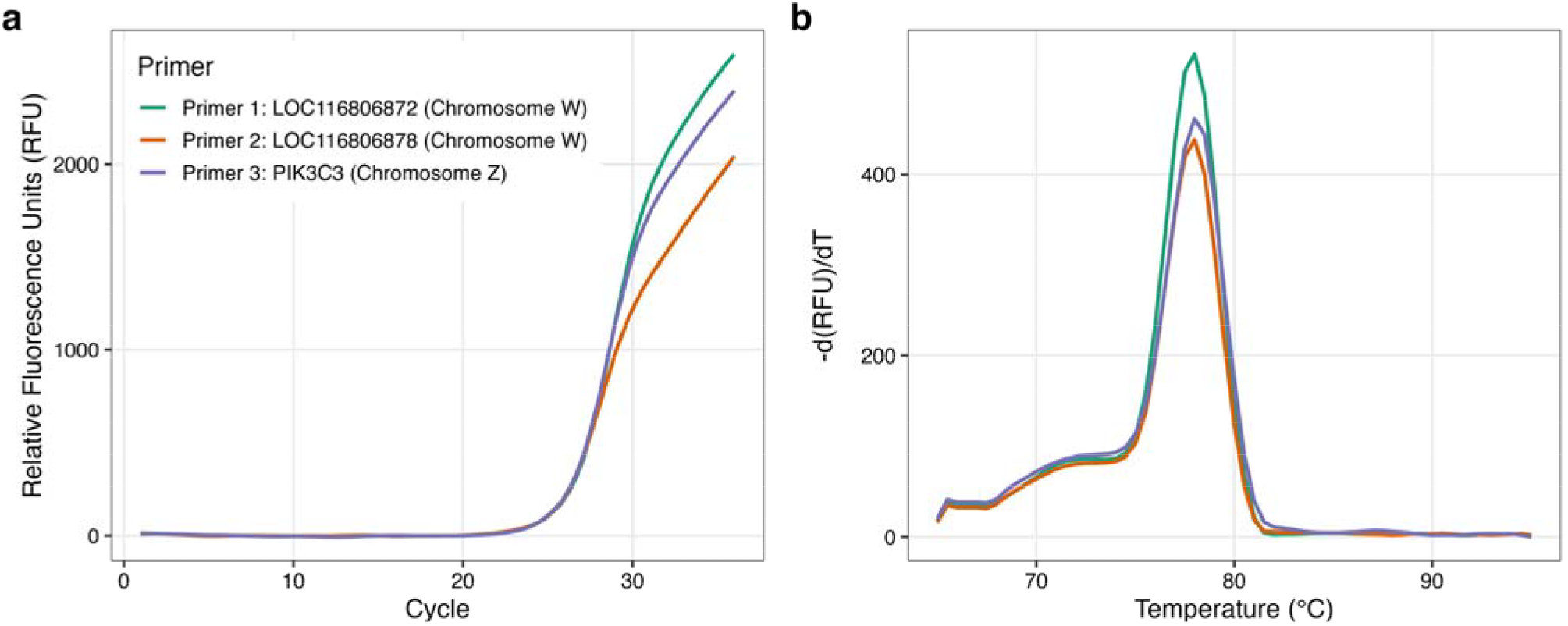
(a) DNA amplification and (b) melting curve peak from a known ZW adult zebra finch female, showing amplification of all three genes.

When applied to E13 zebra finch embryos (2-5ul blood sampled for each), this qPCR assay distinguished individuals as male (ZZ) or female (ZW). RNA sequencing of whole brains of the same individuals supported the sex determined by genomic qPCR. Principal components analysis of the whole brain transcriptome revealed discrete clustering of male and female samples (PC1, 67% of variance in expression, Fig 3a), while quantification of gene expression levels of the markers used in this study confirmed non-overlapping patterns of expression between presumed males and females (Figure 3b-d).

**Figure 3:**
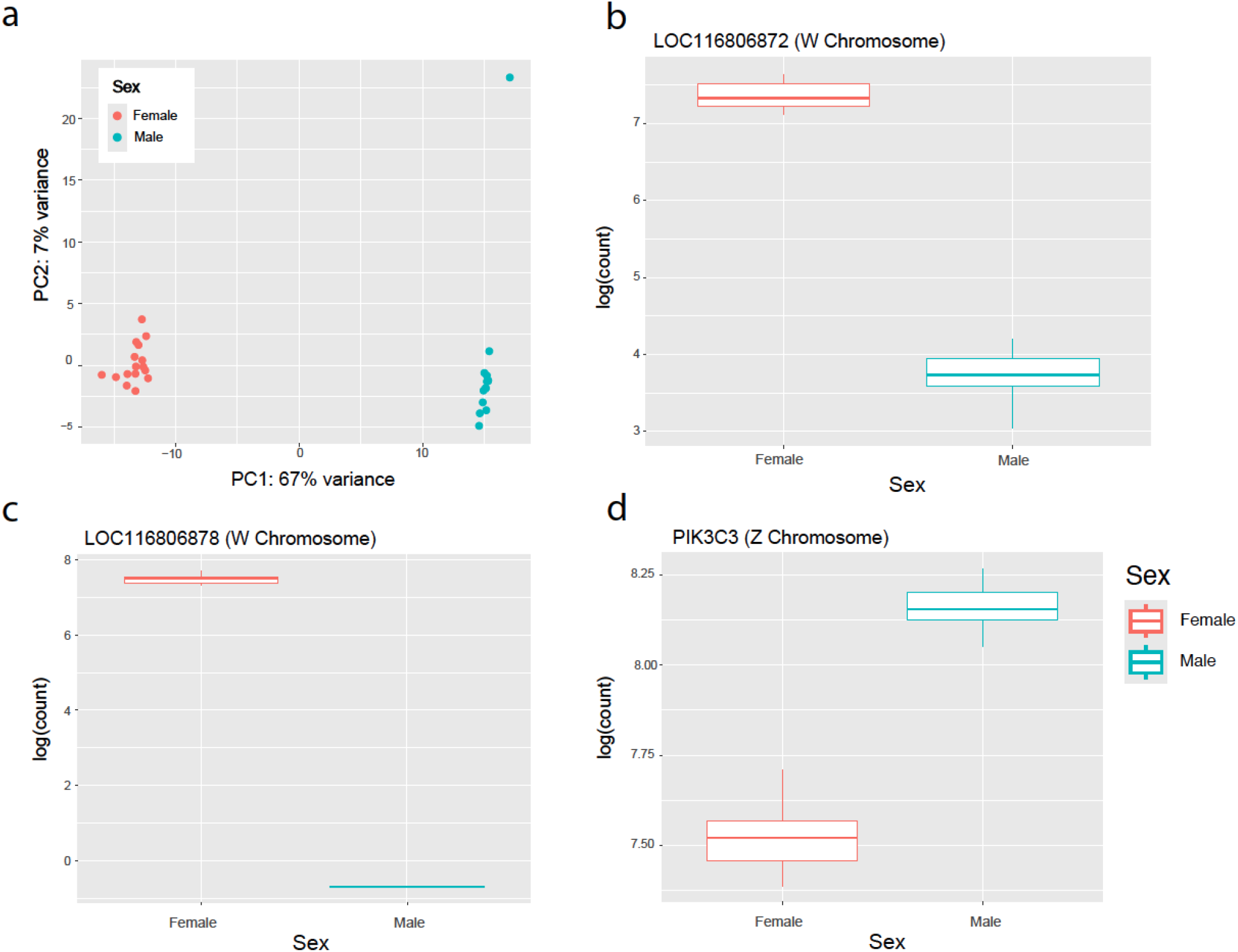
RNA-Sequencing Validation of Gene Expression Differences in Male and Female Zebra Finch Embryos. (a) Principal Component Analysis showing distinct clustering of male and females zebra finch embryos identified initially using our qPCR protocol. Quantification (logarithmic counts) of (b) LOC116806872 (W chromosome), LOC116806878 (W Chromosome), and (d) PIK3C3 (Z chromosome) transcripts in brains of zebra finch embryos.

To assess the cross-species utility of this multi-marker approach, we assayed blood samples from adult chickens of known sex with primers designed for chicken genomic sequences (Table 2), targeting PIK3C3, Nipped-B homolog-like, and spindlin 1 like (SPINL1) genes, the latter having been previously used as a marker for molecular sex identification in chicken (Itoh et al., 2001). The Z target (PIK3C3) was detected in both males and females by qPCR, while the W markers were detected only in females, as predicted (Figures 4 and 5).

**Figure 4:**
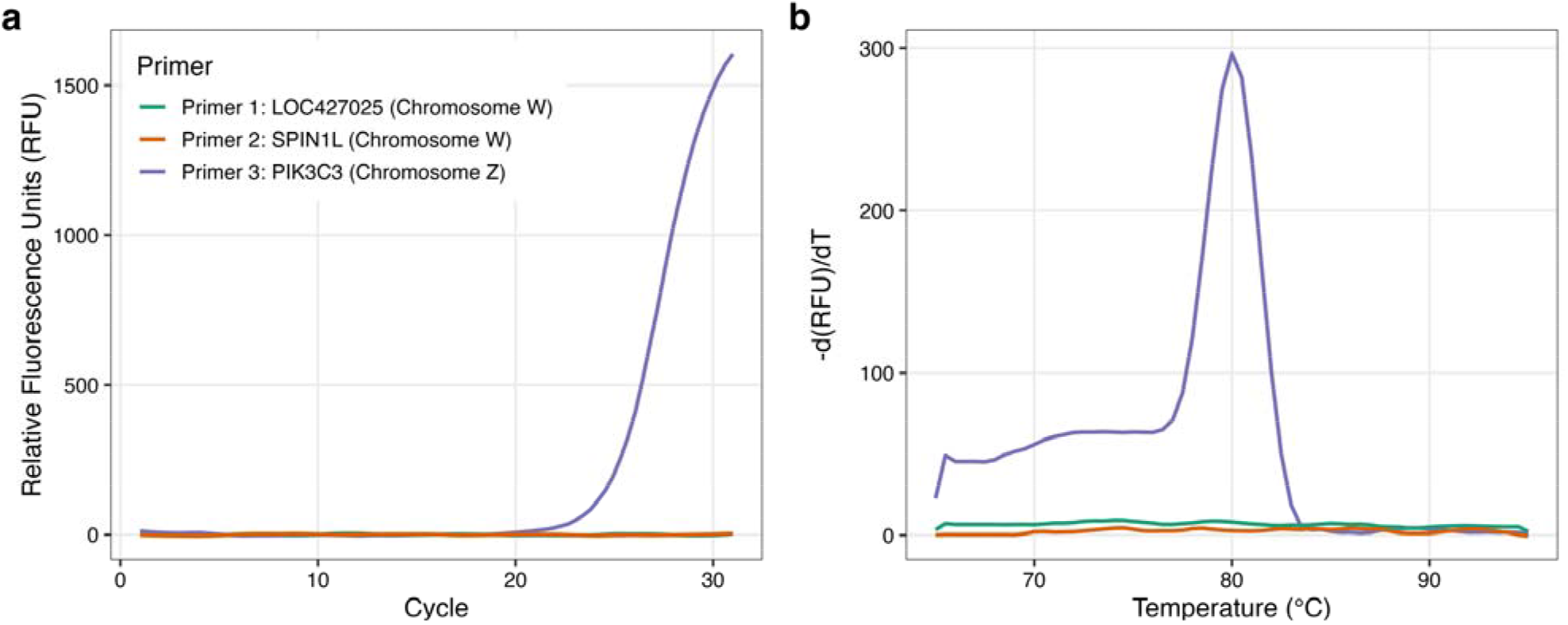
(a) DNA amplification (left) and (b) melting curve peak (right) from a known ZZ adult chicken male produces a single amplification curve for the Z chromosome-specific PIK3C3 gene.

**Figure 5:**
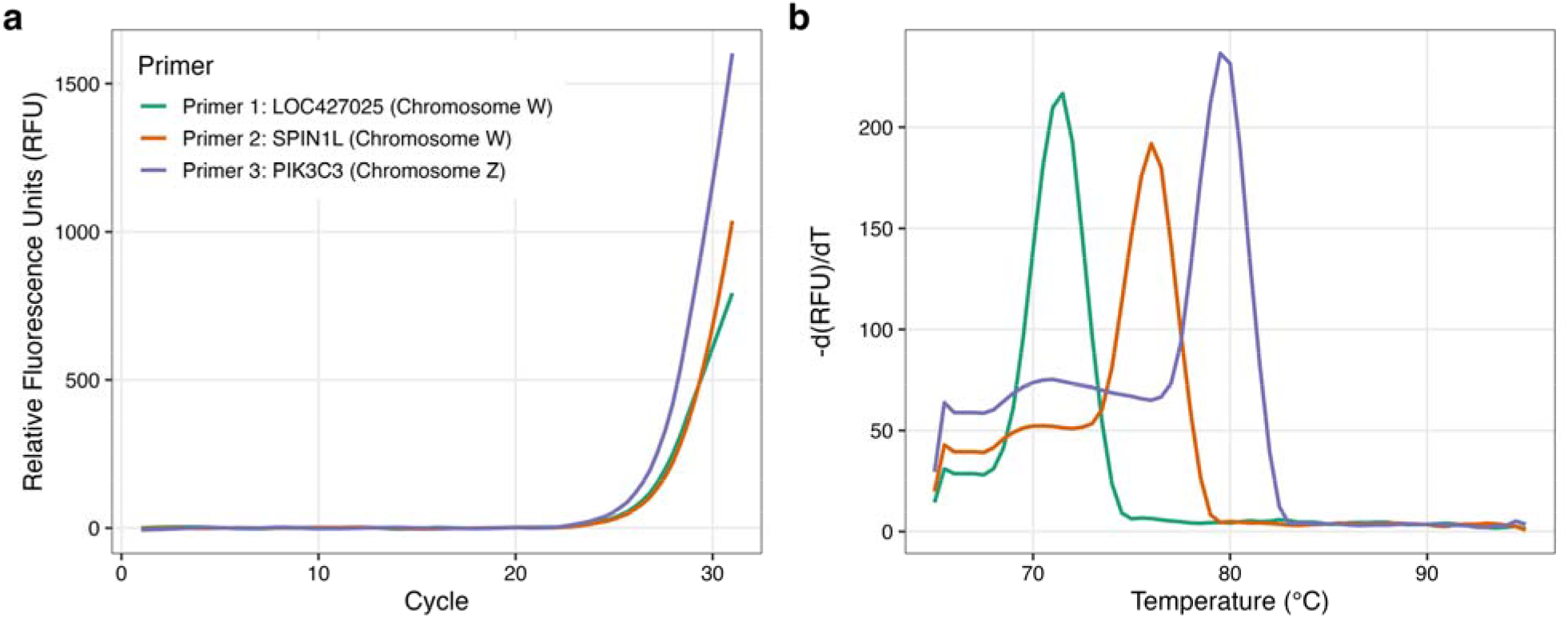
(a) DNA amplification and (b) melting curve peak from a known ZW adult chicken female, showing amplification of all three genes.

## Discussion

Early methods for molecular sexing of birds originated with discovery of the first avian W-encoded gene, CHD1W, which could be detected specifically in females via Southern blot [13]. The subsequent discovery [16] that CHD1W is highly homologous to a Z-encoded gene, CHD1Z, led to the development of PCR assays for genetic sexing that use conserved primers to amplify regions of these two genes that yield different sized PCR products [14,15,17,18], such that males (ZZ) yield a single-sized PCR product, while females (ZW) yield two different-sized PCR products. More recently, this protocol was adapted for real-time qPCR to eliminate the necessity of gel electrophoresis to discriminate the different PCR products [30]. However, this approach can fail due to polymorphism in the CHD1 genes. For example, different CHD1Z alleles in the same male may yield two different-sized PCR products, leading to false assignment of the individual as female [19]. Conversely, the failure of PCR primers to recognize and to amplify CHD1W may lead to false assignment of a female as male [20].

Here we describe a refined assay for determining genetic sex in zebra finch and chickens, which utilizes a SYBR green real-time qPCR assay and newly designed primers targeting two W chromosome markers and one Z marker for each species. The use of SYBR green dye to detect the products of qPCR, followed by melt curve analysis of these products, allows one to ensure that each primer set yields a single product and to read the sexing results directly from the qPCR instrument, without the added effort of separating and visualizing the products by gel electrophoresis. The addition of a second W marker to the assay ensures that failure of amplification by a single primer set (e.g. due to genetic polymorphism) does not lead to false assignment of a sample as male. This assay requires only 1 ng genomic DNA per reaction, which is easily extracted from 2-5 uL of blood. While a commercial gDNA purification kit did yield DNA of higher purity than extraction with Chelex-100 resin, both methods gave similar DNA yield, and both were adequate for the qPCR sexing assay described here.

## Acknowledgements

We acknowledge Allison Rees and Jonathan Roberts for their technical assistance in embryo collection and performing qPCR. We also acknowledge the assistance of Maslyn Greene of the Clemson University Genomics and Bioinformatics Facility (CUGBF) for services and facilities provided for RNA-sequencing. The CUGBF is supported by the College of Science and Grants P20GM146584 and P20GM139769 Institutional Development Awards (IDeA) from the National Institute of General Medical Sciences of the National Institutes of Health.

## References

1. Mello CV. The Zebra finch, taeniopygia guttata: An avian model for investigating the neurobiological basis of vocal learning. Cold Spring Harb Protoc. 2014;2014: 1237–1242.

2. Alvarez-Buylla A, Kirn JR, Nottebohm F. Birth of projection neurons in adult avian brain may be related to perceptual or motor learning. Science. 1990;249: 1444–1446.

3. Schlinger BA, Arnold AP. Circulating estrogens in a male songbird originate in the brain. Proc Natl Acad Sci U S A. 1992;89: 7650–7653.

4. Tagirov M, Rutkowska J. Sexual dimorphism in the early embryogenesis in zebra finches. PLoS One. 2014;9: e114625.

5. Morinha F, Cabral JA, Bastos E. Molecular sexing of birds: A comparative review of polymerase chain reaction (PCR)-based methods. Theriogenology. 2012;78: 703–714.

6. Grüebler MU, Schuler H, Müller M, Spaar R, Horch P, Naef-Daenzer B. Female biased mortality caused by anthropogenic nest loss contributes to population decline and adult sex ratio of a meadow bird. Biol Conserv. 2008;141: 3040–3049.

7. Wheeler ME, Barzen JA, Crimmins SM, Van Deelen TR. Population responses to harvest depend on harvest intensity, demographics, and mate replacement in sandhill cranes. Glob Ecol Conserv. 2021;30: e01778.

8. Mariette MM, Buchanan KL. Prenatal acoustic communication programs offspring for high posthatching temperatures in a songbird. Science. 2016;353: 812–814.

9. Ruiz-Raya F, Noguera JC, Velando A. Covariation between glucocorticoid levels and receptor expression modulates embryo development and postnatal phenotypes in gulls. Horm Behav. 2023;149: 105316.

10. Mariette. Acoustic developmental programming: a mechanistic and evolutionary framework. Trends Ecol Evol. 2021. doi:10.1016/j.tree.2021.04.007

11. Tyson CW, Jennings SL, Hoover BA, Miles A. Energetic constraints drive sex-specific parental care in the monomorphic Leach’s storm-petrelHydrobates leucorhous. J Avian Biol. 2022;2022. doi:10.1111/jav.02904

12. Noguera JC, Velando A. Bird embryos perceive vibratory cues of predation risk from clutch mates. Nat Ecol Evol. 2019. doi:10.1038/s41559-019-0929-8

13. Ellegren H. First gene on the avian W chromosome (CHD) provides a tag for universal sexing of non-ratite birds. Proc Biol Sci. 1996;263: 1635–1641.

14. Griffiths R, Double MC, Orr K, Dawson RJG. A DNA test to sex most birds. Mol Ecol. 1998;7: 1071–1075.

15. Fridolfsson A-K, Ellegren H. A simple and universal method for molecular sexing of non-ratite birds. J Avian Biol. 1999;30: 116.

16. Griffiths R, Korn RM. A CHD1 gene is Z chromosome linked in the chicken Gallus domesticus. Gene. 1997;197: 225–229.

17. Kahn NW, John J, Quinn TW. Chromosome-specific intron size differences in the avian CHD gene provide an efficient method for sex identification in birds. Auk. 1998;115: 1074–1078.

18. Soderstrom K, Qin W, Leggett MH. A minimally invasive procedure for sexing young zebra finches. J Neurosci Methods. 2007;164: 116–119.

19. Casey AE, Jones KL, Sandercock BK, Wisely SM. Heteroduplex molecules cause sexing errors in a standard molecular protocol for avian sexing. Mol Ecol Resour. 2009;9: 61–65.

20. Robertson BC, Gemmell NJ. PCR-based sexing in conservation biology: Wrong answers from an accurate methodology? Conserv Genet. 2006;7: 267–271.

21. Murray JR, Varian-Ramos CW, Welch ZS, Saha MS. Embryological staging of the Zebra Finch, Taeniopygia guttata. J Morphol. 2013;274: 1090–1110.

22. Singh UA, Kumari M, Iyengar S. Method for improving the quality of genomic DNA obtained from minute quantities of tissue and blood samples using Chelex 100 resin. Biol Proced Online. 2018;20: 12.

23. Kraft F-LH, Driscoll SC, Buchanan KL, Crino OL. Developmental stress reduces body condition across avian life-history stages: A comparison of quantitative magnetic resonance data and condition indices. Gen Comp Endocrinol. 2019;272: 33–41.

24. Kraft F-LH, Crino OL, Buchanan KL. Developmental conditions have intergenerational effects on corticosterone levels in a passerine. Horm Behav. 2021;134: 105023.

25. Kraft F-LH, Crino OL, Adeniran-Obey SO, Moraney RA, Clayton DF, George JM, et al. Parental developmental experience affects vocal learning in offspring. Sci Rep. 2024;14: 13787.

26. Qamar W, Khan MR, Arafah A. Optimization of conditions to extract high quality DNA for PCR analysis from whole blood using SDS-proteinase K method. Saudi J Biol Sci. 2017;24: 1465–1469.

27. Bolger AM, Lohse M, Usadel B. Trimmomatic: a flexible trimmer for Illumina sequence data. Bioinformatics. 2014;30: 2114–2120.

28. Patro R, Duggal G, Love MI, Irizarry RA, Kingsford C. Salmon provides fast and bias-aware quantification of transcript expression. Nat Methods. 2017;14: 417–419.

29. Love MI, Huber W, Anders S. Moderated estimation of fold change and dispersion for RNA-seq data with DESeq2. Genome Biol. 2014;15: 550.

30. Brubaker JL, Karouna-renier NK, Chen YU, Jenko K, Sprague DT, Henry PFP. A noninvasive, direct real - time PCR method for sex determination in multiple avian species. Mol Ecol Resour. 2011;11: 415 –417.

